# Long-range regulatory target prediction reveals shared genetic background across ulcerative colitis, Crohn’s disease, primary sclerosing cholangitis and ankylosing spondylitis

**DOI:** 10.64898/2026.06.29.735270

**Authors:** Dorian Dulčić, Katarina Mandić, Dalibor Hršak, Anja Barešić

## Abstract

Common variants detected by the genome-wide association studies (GWAS) create a wealth of knowledge on genetic component of individual traits and diseases. Elucidating the molecular mechanism behind the vast majority of these variants that are found to be non-coding remains a largely unsolved task, especially when distal and pleiotropic interactions between regulatory elements where these variants occur and gene promoters are taken into account. Focusing on four diseases with immune-mediated mechanisms namely ulcerative colitis, Crohn’s disease, primary sclerosing cholangitis and ankylosing spondylitis, we demonstrate the utility of the targPred tool, providing prediction of genes targeted by the regulatory variants. We demonstrate that taking into account evolutionary and comparative genomic data, previously unobserved mechanistic trends (the platelet, vascular and sterol clusters) can be detected in terms of implicated genes targeted by the regulatory elements containing common variants, shared between all four diseases, as well as specific trends for subsets of diseases, e.g. two IBD phenotypes. We also elucidate a clinically-relevant target COG6 shared between IBD and PSC, as well as a whole range of other target genes missed by the conventional SNP-to-gene assignments methods.

## 1 Introduction

Genome-wide association studies (GWAS) have established themselves as one of the principal tools in the discipline of human genetics, cataloguing large amounts of robust statistical associations between common genetic variants and a vast range of traits and diseases [1]. Yet the overwhelming majority of these associations do not map to protein-coding sequence, but to the non-coding passages in the genome: more than 90% of trait-associated variants lie outside coding regions, and they are concentrated in distal regulatory DNA [2]. A statistically associated variant is therefore in most cases a signpost of a region containing regulatory element(s), and identifying the gene through which it exerts its effect is a separate and considerably more difficult task. The default heuristic of assigning a locus to its nearest gene is convenient but frequently misleading, because a regulatory variant may act on a gene hundreds of kilobases away while “skipping over” the interspersed genes. Resolving this gap is a prerequisite for converting association signals into mechanistic insight, accounting for the heritability that remains unexplained by coding variation, and nominating credible therapeutic targets from association data.

Much of the difficulty stems from long-range transcriptional regulation and the inherent cost of experimentally validating chromatin contacts in situ. Enhancers commonly interact with their target promoters across large genomic distances, brought into physical proximity by three-dimensional chromatin folding and looping within a cell, rather than by linear adjacency [3, 4]. Consequently, the regulatory landscape of a single gene may span megabases, organised within the self-interacting chromatin domains that chromosome conformation capture has made visible [5]. A growing family of integrative SNP-to-gene methods now combines such three-dimensional contact information with functional chromatin maps and quantitative-trait data to link variants to genes [6–10], and these approaches have markedly improved locus interpretation. Most of them, however, remain locus-centric and are trained on curated golden standard reference datasets that are necessarily limited in size and biased toward well-studied loci and traits. They also typically operate within a single species, which under-represents both the evolutionary conservation of regulatory wiring and the architectural constraints that enable long-range transcriptional control [11].

An approach grounded in the extreme evolutionary conservation and explicitly oriented toward long-range interactions, is therefore an attractive complement to these widely adopted methods. Genomic regulatory blocks (GRBs) provide exactly such a perspective. GRBs are relatively large genomic regions distinguished by an unexpectedly high density of evolutionarily conserved non-coding elements [12, 13] that surround key developmental genes and encompass their extended, often megabase-scale regulatory landscapes [14]. Within a GRB the distal, often conserved regulatory elements act through chromatin looping bringing them into proximity of the target promoter [3,4]. In this way, one or a few *target* genes are placed under control of these distal elements, while the remaining *bystander* genes are largely unresponsive to them [12]. The genome folds into self-interacting domains detectable by chromosome conformation capture [5], and GRBs generally coincide with individual topologically associating domains, consistent with a shared basis of evolutionary conservation for both phenomena [15]. Because GRB boundaries are defined by conservation across vertebrate evolution rather than by trait-specific data, and because the underlying enhancer-promoter logic is largely invariant across tissues, GRBs offer a principled and economical scaffold for assigning non-coding variants to their long-range targets. This framework has previously been applied to type 2 diabetes and obesity [16] and to neuropsychiatric disorders with specific focus on schizophrenia [17]. Moreover, we recently generalised the GRB approach across the entire GWAS Catalog in the targPred resource [11]. targPred assigns each GWAS locus located within a GRB to its predicted long-range target gene (not by proximity, but on the basis of enhancer-promoter transcriptional activity associations quantified from FANTOM5 expression data across hundreds of biosamples), and benchmarks these predictions against experimentally validated SNP-to-gene sets and other enhancer-gene predictors [11]. For any given trait, targPred thus yields a set of candidate target genes, grounded in regulatory rather than positional evidence, that may diverge from the conventionally reported putative target gene, often heavily guided by the genomic distance.

We apply this long-range perspective to four immune-mediated diseases, chosen for their well-documented genetic and clinical overlap: ulcerative colitis (UC), Crohn’s disease (CD), primary sclerosing cholangitis (PSC) and ankylosing spondylitis (AS). UC and CD are the two principal forms of inflammatory bowel disease (IBD) and share the great majority of their susceptibility loci [18]. PSC, a progressive cholestatic disease of the bile ducts, is strongly comorbid with IBD (roughly three-quarters of PSC patients also suffer from at least early (i.e. subclinical) stage of IBD [19]), yet its genetic architecture is only partly shared with classical IBD [20]. AS belongs to the spondyloarthritis spectrum, which is itself epidemiologically and genetically connected to IBD through shared gut-joint inflammatory mechanisms [21]. Indeed, a systematic cross-disease analysis of five chronic inflammatory conditions encompassing all four diseases studied here, identified numerous susceptibility loci shared across this set of states and concluded that the strong PSC–IBD co-morbidity is best explained by biological pleiotropy rather than by diagnostic overlap [22].

Expectedly similar to other traits and diseases, substantial part of the heritability of these diseases lies in regulatory sequence, and many of the shared variants are themselves non-coding. This makes the quartet an especially informative test case for a regulatory, long-range approach to variant interpretation: if the shared biology of these diseases is encoded, even in part, in distal regulatory relationships, then a GRB-based assignment should be able to expose it where nearest-gene assignment simply cannot. Our central hypothesis is that the genes that differ from the conventional GWAS established assignment (the GRB*̸*=GWAS set) carry their own disease-relevant phenotypic signal, one that complements rather than merely duplicates the signal of the conventionally reported genes, and that this additional regulatory layer aware of interactions over long genomic distances can help to illuminate the biology shared across these four diseases, and in particular the overlap between inflammatory bowel disease and primary sclerosing cholangitis.

## 2 Materials and methods

The primary input to this research was a catalogue of all GWAS loci falling within genomic regulatory blocks (GRBs), where each GWAS locus was linked to its mapped traits and to the gene predicted to be its regulatory target with long-range regulation (LRR) sensitive methodology, as previously implemented in the targPred resource. All genomic coordinates were aligned to the GRCh37/hg19 reference, consistent with the inherited annotations. While no additional processing was performed in Sections 2.1-2.3 when compared to our previous work ([11] and accompanying github page), we also provide a brief overview of this methodology for reader’s convenience.

### 2.1 GRB annotation and target gene prediction

GRB intervals (based on the human:dog non-coding conservation) and associated target gene assignments (based on the random forest model of key attributes discerning target from bystander genes) were obtained from the targPred, with no additional processing.

### 2.2 GWAS variants and genomic loci

In the targPred resource, a GWAS locus is defined as the linkage-disequilibrium (LD) block surrounding a tagging reported SNP. It is worth noting here that LD blocks were computed on the 1000 Genomes phase 3 (GRCh37) reference panel restricted to the European superpopulation [23], and only GWAS studies of European-ancestry participants were included.

### 2.3 GWAS loci in GRBs and enhancer-promoter associations

As utilised previously, locus is assigned to a GRB when it shares any nucleotide overlap with the GRB interval [11, 17] and only loci colocalising with a GRB are reported here unless otherwise stated. The target gene assignment for each GRB are not made by proximity, but rather constructed based on functional effect. Briefly, for every enhancer-promoter pair within a GRB, the association between enhancer transcriptional activity and gene expression is quantified as an empirical *p*-value (empP) from FANTOM5 CAGE data across 309 biosamples encompassing a remarkable range of human cell and tissue contexts and developmental stages, capturing both activating and repressive directions of the regulation.

### 2.4 Gene set classification and agreement

From the initial dataset of variants, LD blocks, GWAS target gene predictions, targPred (GRB-based) target gene predictions, and enhancer-promoter activity associations, we restricted the locus selection to four diseases of interest: “ulcerative colitis”, “Crohn’s disease”, “primary sclerosing cholangitis” and “ankylosing spondylitis”. This was done by applying filtering on the “mapped trait” column for the exact disease terms, further retaining only associations reaching genome-wide significance (*P <* 5×10^−8^) [1] using a locally created R script. No GWAS summary statistics were downloaded and no loci were defined in this study; the underlying associations and loci originate from the GWAS Catalog (https://www.ebi.ac.uk/gwas/) as processed within the targPred resource. The complete list of loci, with disease membership by locus, is provided in Table S1.

To further facilitate biological interpretation of obtained results, each gene was classified into one of three mutually exclusive categories. The first, denoted GRB=GWAS (*n* = 13), comprises genes whose targPred target-gene assignment coincides with the gene reported or mapped for the same locus in the GWAS Catalog (i.e. LRR-sensitive prediction agrees with the conventionally assigned gene). The second, GRB*̸*=GWAS (*n* = 61), comprises genes predicted as GRB targets that differ from the GWAS reported gene for the given locus. The third, designated secondary (*n* = 12), comprises genes showing GWAS overlap in two or three of the four diseases, treated separately because of the valuable insight into the differential cross-disease signal among the diseases of interest.

## 3 Results

The analysis presented below starts with all GWAS variants associated with the four diseases of interest, namely ulcerative colitis, Crohn’s disease, primary sclerosing cholangitis and ankylosing spondylitis. The full list of these SNPs is provided as Table S2, with accompanying GRB localisation status: in GRB or out of a GRB. The latter ones are hereafter eliminated as these are not presumed to be involved in LRR and therefore fall beyond the scope of this study.

### 3.1 Candidate gene catalogue and classification

Our first focus was on predicted target genes by the targPred method, as potential expansion of the GWAS-mapped target genes for the non-coding variants, often neglecting long-range promoter-enhancer effects. To that end, we focused on the statistically significant genome-wide associations (*P <* 5×10^−8^) between loci associated with 4 diseases and genes as calculated by the targPred method. This yielded 145 colocalised loci associated with genes in genomic regulatory blocks, revealing 86 associated GRB target genes putatively under LRR for these phenotypes. Classifying each gene by whether its targPred assignment coincides with the gene reported or mapped for the locus in the GWAS Catalog (typically the nearest or locally prioritised gene) shows that only 13 of the 86 genes (15%) agree with the conventional assignment (GRB=GWAS), whereas 61 (71%) have targPred-inferred targets that differ from it (GRB*̸*=GWAS). A further 12 (14%) recur across only two or three of the four diseases, marking a separate category (termed secondary) of particular biological interest. For the majority of the disease loci, in other words, the gene associated by taking into account long-range regulatory evidence is not the one a conventional GWAS approach would report, which is the key observation that both motivates and frames the phenotype analyses that follow.

The cross-disease structure of the data, or how the four diseases intersect at successive levels of resolution, is summarised in the panels of Figure 1, which present the overlaps across UC, CD, PSC and AS of, respectively, the associated SNPs, the GWAS-reported target genes, the targPred reported target genes, and the underlying loci. A subset of loci returned missing (NA) target-gene assignments for one or more of the four diseases in the automated targPred output; each of these was resolved individually by manual lookup and curation before the candidate set was finalised. Because the four panels of Figure 1 are computed from the automated per-disease assignments, the targPred target-gene panel (Figure 1D) enumerates fewer genes than the 86 distinct candidates obtained after this curation. The secondary set defined above corresponds to the multi-disease intersections of the targPred target-gene panel. The complete catalogue of these 86 candidate genes, with their category, cross-disease overlap and a representative locus, is given in Table 1.

**Figure 1:**
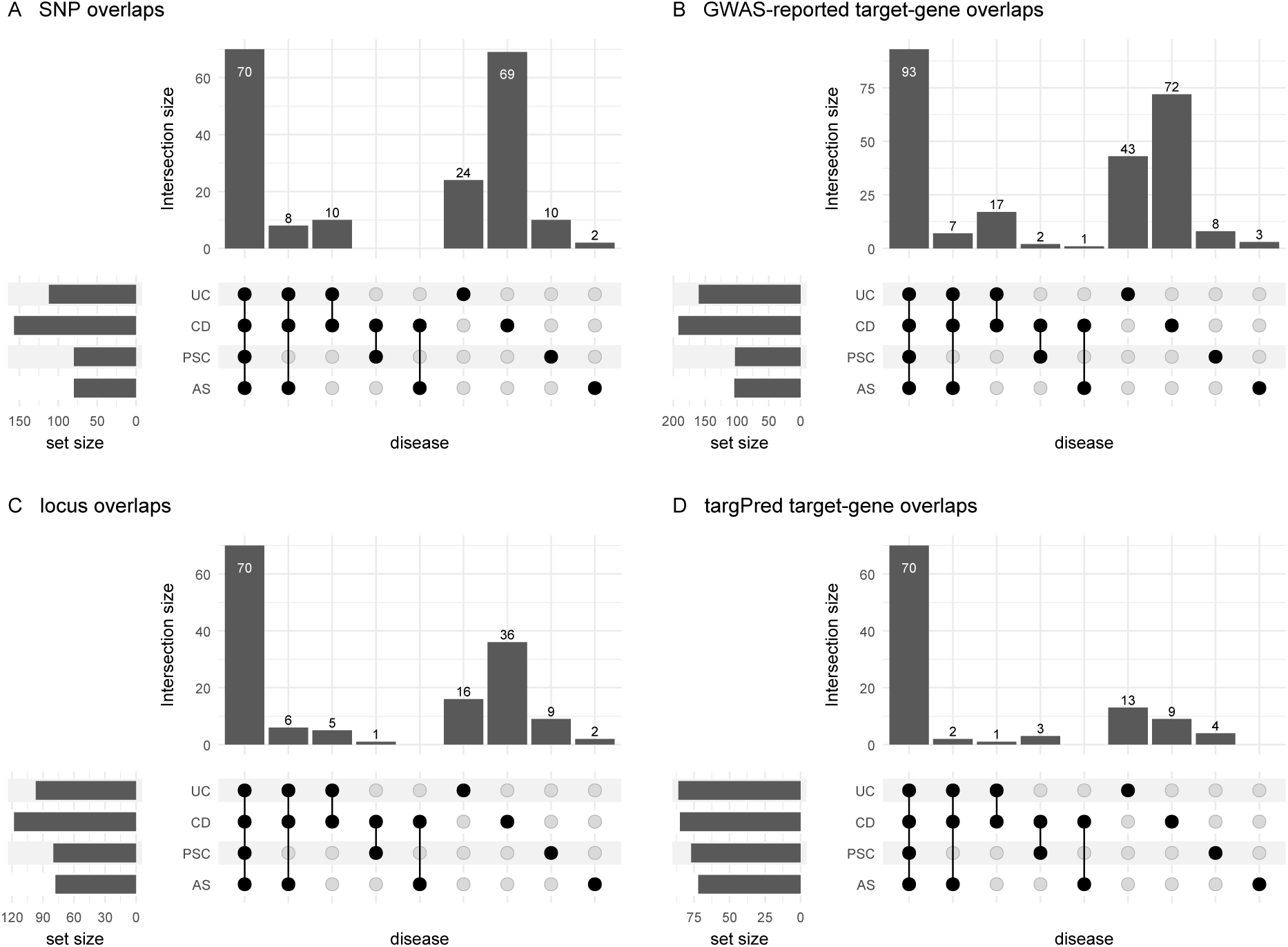
Cross-disease overlap across UC, CD, PSC and AS at four levels of resolution. (A) Number of associated SNPs, (B) Number of GWAS-reported target genes, (C) Number of the underlying loci and (D) Number of targPred target genes. The secondary gene set corresponds to the two- and three-disease intersections of panel D.

**Table 1:** The 86 candidate GRB target genes for UC, CD, PSC and AS, by category. The **GWAS gene** column gives the gene reported or mapped for the locus in the GWAS Catalog: for GRB=GWAS it coincides with the targPred-inferred target; for GRB*̸*=GWAS the differing reported gene is shown (– where the only reported gene is the target itself). The twelve genes recurring across two or three diseases (the secondary set) are placed in GRB=GWAS or GRB*̸*=GWAS according to that locus. For genes mapping to several loci the lead locus is shown with the number of additional loci in parentheses; full loci and SNPs are in the accompanying CSV.

| targPred target | GWAS gene | Diseases | Locus (lead SNP) |
| --- | --- | --- | --- |
| **GRB=GWAS** ($n = 18$) | | | |
| CD28 | CD28 | all 4 | chr2:204585502-204649276 – rs7426056 |
| FAM53B | FAM53B | UC, CD | chr10:126316604-126551228 – rs111456533 |
| FOS | FOS | all 4 | chr14:75702235-75747118 – rs1569328 |
| FYN | FYN | UC, CD | chr6:111493953-111916163 – rs3851228 |
| IL21 | IL21 | all 4 | chr4:123008516-123555178(+3) – rs13140464 |
| JAZF1 | JAZF1 | all 4 | chr7:28142088-28214282(+1) – rs10276381 |
| LPHN2 | LPHN2 | UC, CD, AS | chr1:82227391-82252004 – rs2066363 |

Table 1 (continued)
| targPred target | GWAS gene | Diseases | Locus (lead SNP) |
| --- | --- | --- | --- |
| NKD1 | NKD1 | all 4 | chr16:50627378-50694011(+1) – rs117372389 |
| NKX2-3 | NKX2-3 | all 4 | chr10:101271789-101295863(+2) – rs10748781 |
| NR5A2 | NR5A2 | all 4 | chr1:200065713-200105746 – rs2816958 |
| PDGFB | PDGFB | all 4 | chr22:39659773-39737810(+1) – rs2143178 |
| PRDM1 | PRDM1 | all 4 | chr6:106435025-106442096(+3) – rs28701841 |
| PTPRK | PTPRK | all 4 | chr6:128215237-128297611(+1) – rs13204742 |
| RUNX3 | RUNX3 | all 4 | chr1:25289734-25305783 – rs6600247 |
| SMAD7 | SMAD7 | UC, CD | chr18:46395022-46395022 – rs7240004 |
| SPRED2 | SPRED2 | all 4 | chr2:65557287-65693513(+1) – rs11675538 |
| TENM3 | TENM3 | UC, CD, AS | chr4:183745321-183748510 – rs7660520 |
| TRIB1 | TRIB1 | all 4 | chr8:126527765-126541090 – rs1551398 |
| **GRB≠GWAS** ($n = 68$) | | | |
| ABCG5 | THADA | all 4 | chr2:43451957-43864089 – rs77981966 |
| ABCG8 | THADA | all 4 | chr2:43451957-43864089 – rs77981966 |
| ARHGAP15 | TEX41 | all 4 | chr2:145417530-145627269 – rs11681525 |
| BRE | FOSL2, FLJ31356 | all 4 | chr2:28602911-28647084 – rs925255 |
| BRINP1 | TLR4 | all 4 | chr9:120435875-120475602 – rs4986790 |
| C1QC | ZBTB40 | all 4 | chr1:22681214-22711473(+1) – rs12568930 |
| CD83 | RNU6-793P, PPIAP9 | all 4 | chr6:14711961-14734463 – rs1267499 |
| CDH22 | CD40, PLTP | all 4 | chr20:44680853-44749251 – rs1883832 |
| CLTC | RPS6KB1 | all 4 | chr17:57800966-58046246(+1) – rs1292035 |
| COG6 | LINC00598 | UC, CD, PSC | chr13:40678443-40706358 – rs915286 |
| COL2A1 | HDAC7, VDR | all 4 | chr12:48195939-48208368 – rs11168249 |
| CPEB4 | – | all 4 | chr5:173269956-173393823(+1) – rs17695092 |
| CUX2 | SH2B3, ATXN2 | all 4 | chr12:111826477-112273499 – rs3184504 |
| CYP26A1 | HHEX | all 4 | chr10:94248310-94495241 – rs2497318 |
| CYP26C1 | HHEX | all 4 | chr10:94248310-94495241 – rs2497318 |
| EGR2 | ZNF365 | all 4 | chr10:64342847-64377503(+5) – rs10761648 |
| ENOX1 | LACC1, CCDC122 | all 4 | chr13:44406102-44490181 – rs3764147 |
| EPHA7 | BACH2 | all 4 | chr6:90656307-90673789(+3) – rs17765610 |
| ETS1 | – | all 4 | chr11:128380974-128396738 – rs11221332 |
| EVX1 | SKAP2 | all 4 | chr7:26694926-26911904(+1) – rs10486483 |
| FLI1 | LINC02098 | all 4 | chr11:128191854-128196675(+1) – rs11221332 |
| FLRT1 | CCDC88B, ESRRA | CD, PSC | chr11:64012910-64141771 – rs663743 |
| FOXD1 | TMEM174 | all 4 | chr5:72515051-72559339 – rs34804116 |
| HCRT | STAT3 | all 4 | chr17:40466092-40545770(+2) – rs12942547 |
| HDGFL1 | CDKAL1 | all 4 | chr6:20623084-20726035(+3) – rs12663356 |
| HEY1 | MIR5708, RNU6-1213P | UC, CD | chr8:81220185-81315490 – rs12543811 |
| HHEX | EIF2S2P3 | all 4 | chr10:94248310-94495241 – rs2497318 |
| HOXA1 | SKAP2 | all 4 | chr7:26694926-26911904(+1) – rs10486483 |
| HOXA10 | SKAP2 | all 4 | chr7:26694926-26911904(+1) – rs10486483 |
| HOXA11 | SKAP2 | all 4 | chr7:26694926-26911904(+1) – rs10486483 |
| HOXA13 | SKAP2 | all 4 | chr7:26694926-26911904(+1) – rs10486483 |
| HOXA3 | SKAP2 | all 4 | chr7:26694926-26911904(+1) – rs10486483 |
| HOXA4 | SKAP2 | all 4 | chr7:26694926-26911904(+1) – rs10486483 |
| HOXA5 | SKAP2 | all 4 | chr7:26694926-26911904(+1) – rs10486483 |
| HOXA6 | SKAP2 | all 4 | chr7:26694926-26911904(+1) – rs10486483 |
| HOXA7 | SKAP2 | all 4 | chr7:26694926-26911904(+1) – rs10486483 |
| HOXA9 | SKAP2 | all 4 | chr7:26694926-26911904(+1) – rs10486483 |
| IKZF1 | SPATA48, C7orf72 | all 4 | chr7:50257634-50323456 – rs1456893 |
| IL20RB | NPM1P17 | all 4 | chr3:137372132-137443557 – rs6781808 |
| KCNH7 | IFIH1 | all 4 | chr2:162992004-163132346(+3) – rs2111485 |
| LHX9 | DENND1B, C1orf53 | all 4 | chr1:197342380-197813558 – rs16841904 |
| LIF | UQCR10 | all 4 | chr22:30130115-30590222(+2) – rs1003342 |
| LIX1 | LNPEP, LRAP | all 4 | chr5:96200770-96373750 – rs1363907 |
| MSX1 | CYTL1 | UC, CD, AS | chr4:4990755-4992367 – rs7672495 |
| NEUROD2 | GSDMB, ORMDL3 | all 4 | chr17:37902887-38089717 – rs12946510 |
| NOG | C17orf67 | all 4 | chr17:54868990-54949047 – rs3853824 |
| NOL4L | POFUT1, KIF3B | all 4 | chr20:30696392-31099311 – rs4243971 |
| NR4A2 | GPD2 | UC, CD | chr2:157596521-157642017 – rs8179646 |
| OLIG3 | SLC17A1 | all 4 | chr6:137959235-138006504 – rs6920220 |
| PLAU | FUT11, CHCHD1 | all 4 | chr10:75469091-75695724 – rs2227551 |
| POLR3A | ZMIZ1 | all 4 | chr10:81032532-81048611(+3) – rs1250544 |
| PRDM8 | ANTXR2 | all 4 | chr4:80816695-80962712 – rs11098964 |
| RBMS3 | – | all 4 | chr3:28215960-28577676 – rs10510607 |
| RFX4 | RIC8B | all 4 | chr12:107052620-107291383 – rs12369214 |

Table 1 (continued)
| targPred target | GWAS gene | Diseases | Locus (lead SNP) |
| --- | --- | --- | --- |
| RP11-574K11.31 | PLAU, FUT11 | all 4 | chr10:75469091-75695724 – rs2227551 |
| SALL1 | NOD2 | all 4 | chr16:50746294-50809536(+2) – rs13333062 |
| SAMSN1 | CYCSP42 | all 4 | chr21:16784168-16838662(+1) – rs1297265 |
| SATB1 | SATB1-AS1 | all 4 | chr3:18610175-18706858(+1) – rs13073817 |
| SATB2 | RFTN2, PLCL1 | all 4 | chr2:198386175-198954831(+5) – rs1016883 |
| SKOR1 | SMAD3 | all 4 | chr15:67441750-67468285(+1) – rs17293632 |
| SLC12A5 | CD40, PLTP | all 4 | chr20:44680853-44749251 – rs1883832 |
| SLC45A1 | TNFRSF9, PARK7 | all 4 | chr1:7969507-8186232 – rs35675666 |
| SPRY4 | NDFIP1 | all 4 | chr5:141435466-141543989(+1) – rs11167764 |
| TUBB1 | ZNF831, CTSZ | all 4 | chr20:57809343-57829821 – rs259964 |
| TYRP1 | LURAP1L | UC, CD, AS | chr9:12785073-12790262 – rs7042370 |
| VEGFB | CCDC88B, ESRRA | CD, PSC | chr11:64012910-64141771 – rs663743 |
| WNT11 | EMSY, C11orf30 | all 4 | chr11:76270683-76302073(+1) – rs11236797 |
| WNT4 | ZBTB40 | all 4 | chr1:22681214-22711473(+1) – rs12568930 |

### 3.2 Shared disease-relevant pathway networks

The candidate genes that recur within shared disease-relevant pathways fall into a small number of mechanistic networks, whose membership across all 86 genes is mapped in Figure 2. The largest cluster marked in red, consists of the lymphocyte master transcription-factors and is shown in greater detail in Figure 3. The next cluster of genes reveals platelet signature (including traits such as increased mean platelet volume, macrothrombocytopenia and impaired platelet aggregation), shown in Figure 4. The following cluster of interest for the potential causal overlap between IBD as a whole and PSC is COG6, shown in Figure 5. Finally, three secondary modules (relevant for subset of the four explored diseases) are shown in Figure 6.

**Figure 2:**
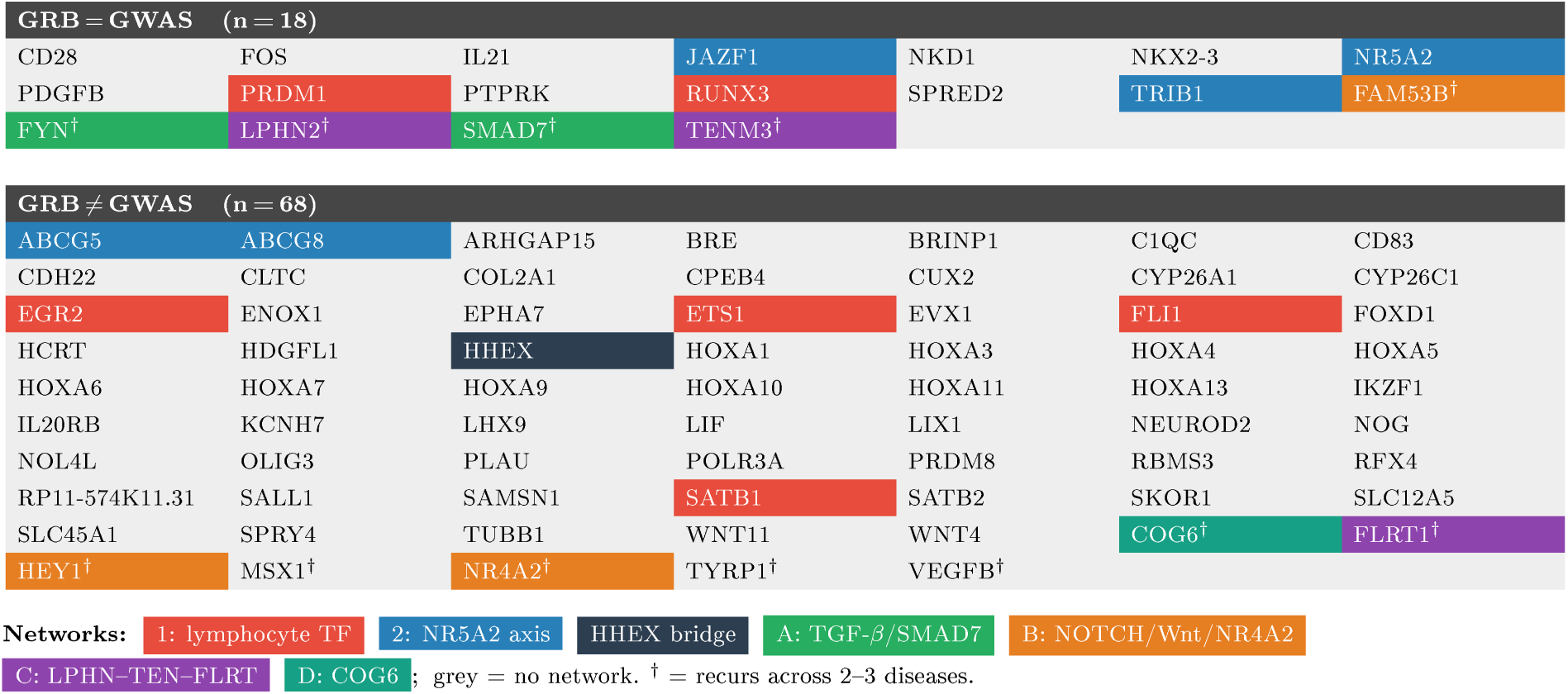
The 86 candidate GRB target genes, classified by agreement with the GWAS-reported gene and coloured by mechanistic-network membership. Genes are split into GRB = GWAS (*n* = 18; the predicted targPred target gene coincides with the GWAS-reported gene at the locus) and GRB *̸*= GWAS (*n* = 68; the target differs between two methods). Fill colour marks the network to which each gene is assigned: Network 1, the lymphocyte master transcription-factor programme (red); Network 2, the NR5A2 gut-liver-pancreas axis (blue); and the secondary modules A (TGF-*β*/SMAD7, green), B (NOTCH/Wnt with NR4A2, orange), C (latrophilin-teneurin-FLRT, purple) and D (COG6 glycosylation, teal). HHEX (dark) is shared between Networks 1 and 2; grey cells are genes not assigned to any network. Genes marked ^†^ recur across two or three of the four diseases (the secondary set) and are placed in their true category rather than kept separate.

**Figure 3:**
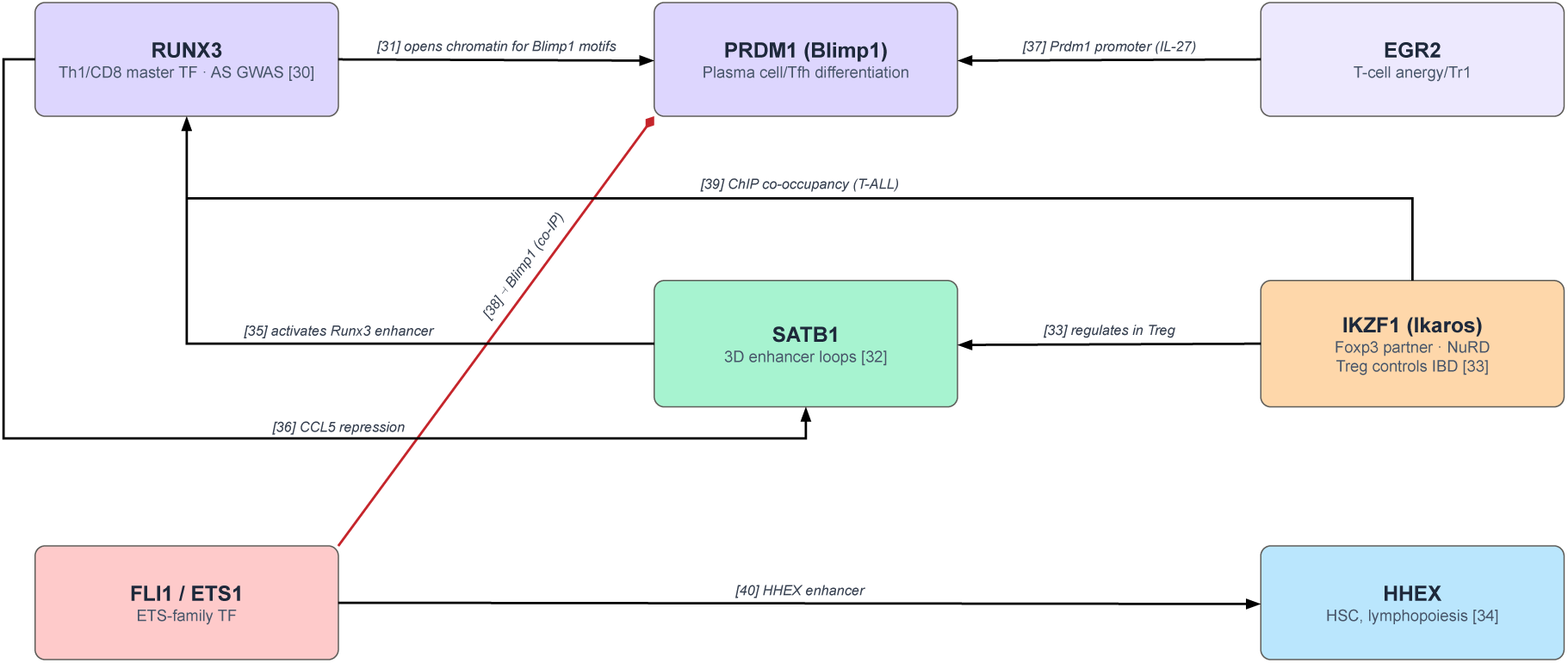
Network 1: lymphocyte master transcription-factor programme. targPred targets RUNX3, PRDM1, SATB1, IKZF1, ETS1, FLI1, HHEX and EGR2 converging on a shared developmental T-cell regulatory module.

**Figure 4:**
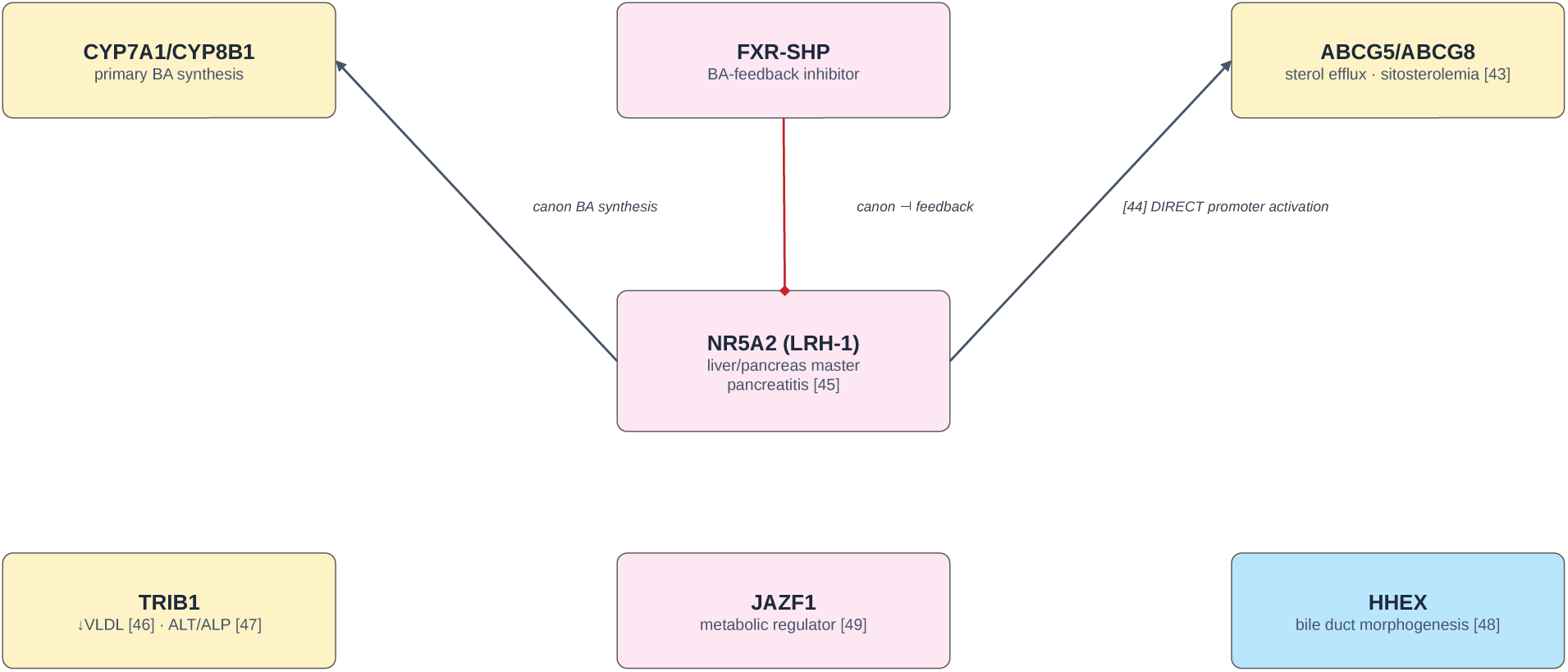
Network 2: NR5A2 nuclear-receptor gut–liver–pancreas axis. NR5A2/LRH-1 with ABCG5, ABCG8, TRIB1, HHEX and JAZF1, of particular relevance to primary sclerosing cholangitis.

**Figure 5:**
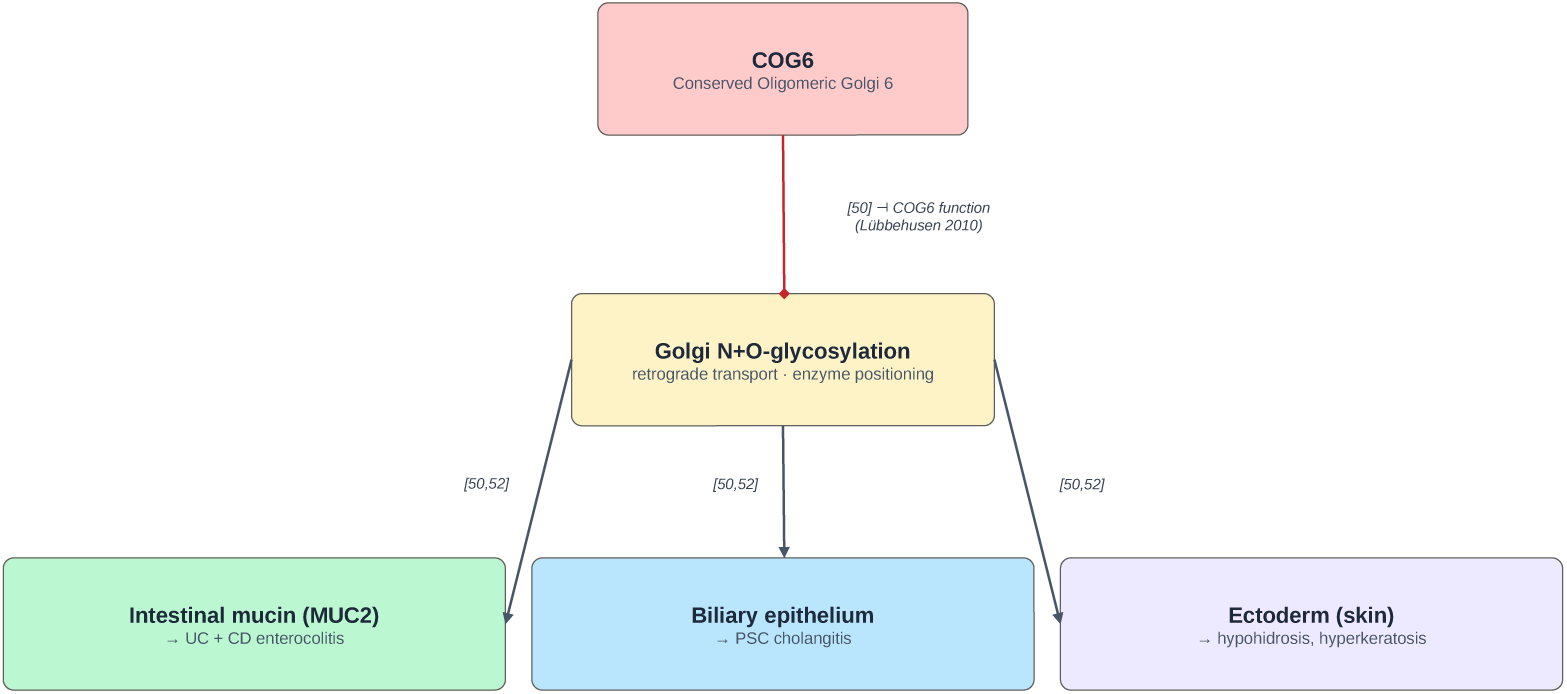
COG6 glycosylation gut–liver axis. The conserved oligomeric Golgi subunit COG6 nominated for the PSC–IBD overlap.

**Figure 6:**
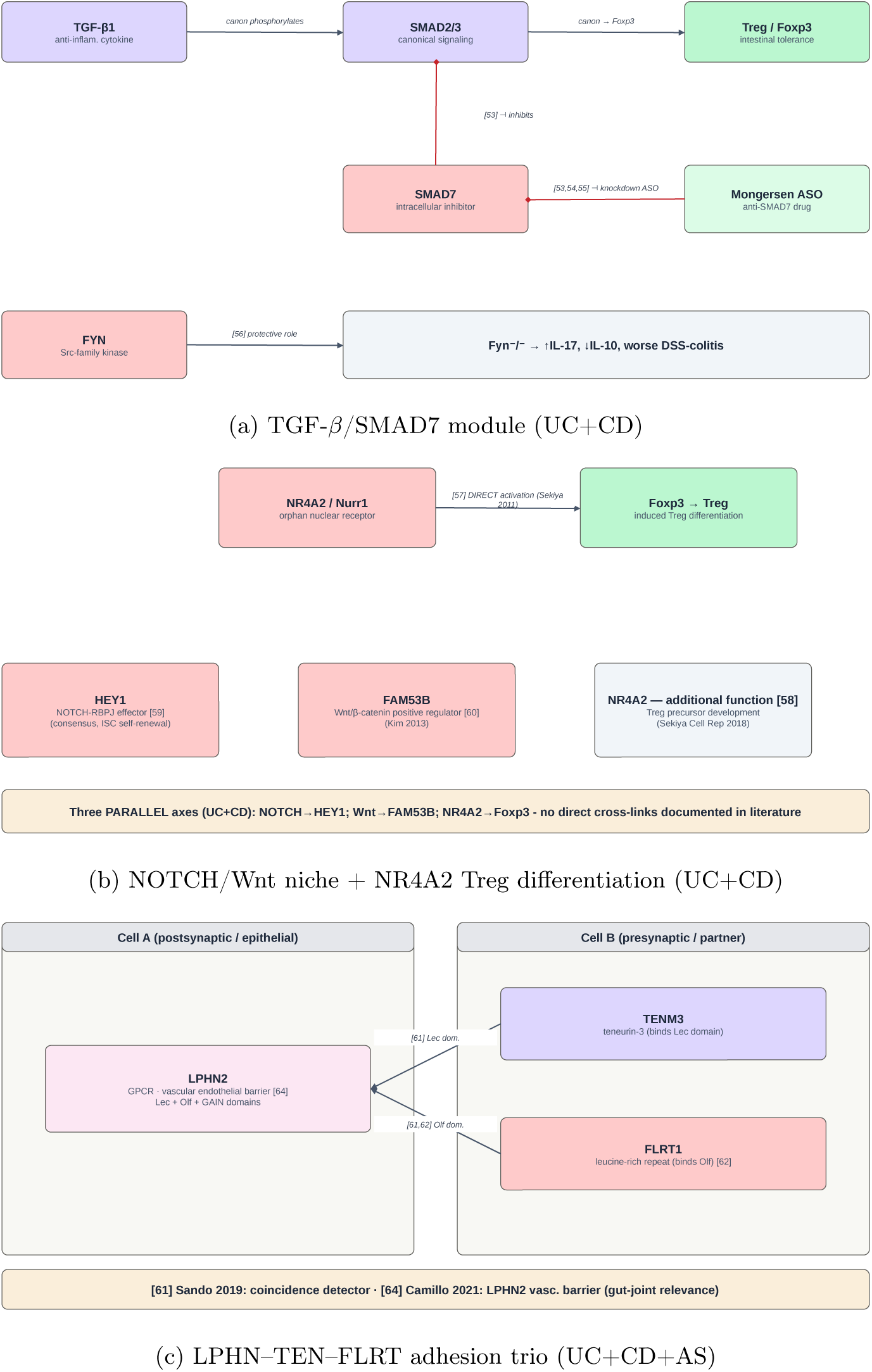
Secondary signalling and adhesion modules. The TGF-*β*/SMAD7 module with FYN, the NOTCH/Wnt niche module around NR4A2 (with HEY1 and FAM53B), and the latrophilin–teneurin–FLRT adhesion trio.

## 4 Discussion

By assigning each disease associated locus from the GWAS catalog to the target gene predicted to be its long-range regulatory target through the targPred framework, rather than to the most proximal or conventionally reported gene, we partitioned the candidate genes into a congruent set (GRB=GWAS), in which the GWAS and targPred assignments agree, and a disjunct set (GRB*̸*=GWAS), in which they diverge. Next, we consider networks presented in Figure 2 in turn.

The immune signature of the congruent set is consistent with a developmentally associated lymphocyte transcription factor programme that recurs among these genes (Figure 3). RUNX3, which is itself associated with ankylosing spondylitis [24], establishes cis-regulatory chromatin accessibility in cytotoxic T cells [25]; SATB1 organises the three-dimensional enhancer network of the developing T-cell genome [26]; IKZF1 acts alongside FOXP3 to control regulatory T-cell gene expression [27]; and HHEX is required for physiologically adequate lymphopoiesis [28]. Beyond their individual roles these factors are wired into a single regulatory circuit: SATB1 helps specify T-lymphocyte subsets and feeds back onto the *Runx3* enhancer [29], RUNX3 in turn antagonises *CCL5* enhancers [30], EGR2 supports Blimp-1 (PRDM1)-dependent IL-10 induction [31], ETS1 restrains Blimp-1 activity during plasma-cell differentiation [32], IKZF1 co-occupies target loci with GFI1 [33], and the ETS-family factors FLI1 and ETS1 converge on the *HHEX* enhancer as part of this shared regulatory circuit [34]. Importantly, this single programme is populated by genes from both categories: RUNX3 coincides with the conventionally reported GWAS gene (GRB=GWAS), whereas SATB1, IKZF1 and HHEX emerge only from the disjunct gene set (GRB*̸*=GWAS). The disjunct set of targets therefore do not diverge from or override the GWAS signal; they complement it, supplying additional, disease-relevant members of the same regulatory module that nearest-gene assignment leaves incomplete. This is consistent with the GRB framework, where such developmental regulators have been demonstrated to often be found under LRR [12, 17].

Beyond complementing this immune programme, the disjunct set also points to biology not represented among the congruent set of genes at all. The second largest network is driven by genes with established roles in megakaryocyte and platelet biology (Figure 4), including the ETS-family transcription factor FLI1, a regulator of megakaryopoiesis whose hemizygous loss underlies the pathogenesis of the Paris-Trousseau macrothrombocytopenia subtype [35], and the platelet-specific *β*1-tubulin TUBB1, mutations of which cause congenital macrothrombocytopenia [36]; notably, platelet dysfunction is itself clinically documented in both IBD and PSC. The vascular and calcification signal converges on the sterol transporters ABCG5 and ABCG8, whose loss causes sitosterolemia [37] and whose promoter is activated by the nuclear receptor NR5A2/LRH-1 [38]. NR5A2 sits at the centre of a gut-liver-pancreas axis of particular relevance to PSC: it is implicated in pancreatic inflammation [39], TRIB1 influences hepatic lipid handling and circulating liver enzymes [40, 41], and HHEX is required for bile-duct morphogenesis [42]; the metabolic regulator JAZF1, implicated in type 2 diabetes, adds a further component to this axis [43]. ABCG5 and ABCG8 bridge the platelet and vascular clusters, linking sterol biology to both (Figure 4).

Next, it is worth noting a shared signature between IBD (i.e. UC and CD) and PSC in the form of COG6 (Figure 5). Deficiency of said gene, which generates a conserved oligomeric subunit of the Golgi apparatus, causes a congenital glycosylation disorder [44, 45] whose phenotype spans disruption of intestinal secretion of mucin (relevant to UC and CD) and disruption of homeostasis of the biliary epithelium (relevant to PSC) [46], mirroring exactly the disease combination of interest. A separate cross-autoimmune cluster (C1QC, ETS1, JAZF1), associated primarily with discoid lupus rash, suggests shared genetic architecture with systemic lupus erythematosus.

Finally, several targPred-inferred targets also fall on therapeutically or mechanistically relevant modules (Figure 6), specific to IBD, or IBD+AS. SMAD7, an intracellular inhibitor of TGF-*β* signalling, is the target of an antisense oligonucleotide named Mongersen, which alleviated Crohn’s disease in early clinical trials [47], before a phase 3 study was halted due to insufficient clinical effect [48], a discordance with the earlier trial that has since been re-examined [49]; the co-occurring FYN is observed to have a protective effect against experimentally induced colitis in animal models [50]. NR4A2 directly induces FOXP3 and shapes regulatory T-cell differentiation [51], with NR4A receptors more broadly governing the development and survival of labile regulatory T-cell precursors [52], and sits alongside two further signalling targets relevant for cellular niches that recur in UC and CD: HEY1, a canonical NOTCH-RBPJ effector [53], and FAM53B, a positive regulator of Wnt/*β*-catenin signalling [54]. The latrophilin-teneurin-FLRT adhesion trio (LPHN2, TENM3, FLRT1), which recurs across UC, CD and AS, forms a coincidence-detection complex [55], the structural basis and synaptic target-recognition role of which have since been resolved [56, 57]; LPHN2 additionally controls endothelial barrier function [58], which represents a plausible point of contact between gut and joint involvement.

These interpretations, however, have certain limitations, the target-gene predictions and the GRB definitions were inherited from the targPred resource [11] and not re-derived here, and the mechanistic networks above are assembled from the literature and are hypothesis-generating rather than causal. Despite extensive benchmarking of the targPred predictions against experimentally validated SNP-to-gene associations and against established enhancer-gene predictors, which established the credibility of the assignments inherited and analysed in this study [11], each predicted regulatory association (both by the GWAS and the targPred method), will require direct functional validation.

Taken together, these observations illustrate the utility of the targPred approach beyond conventional interpretation of disease-associated genetic loci. First, it connected what would otherwise remain disconnected, gathering dispersed disease loci into coherent regulatory modules (most visibly, the platelet, vascular and sterol cluster bridged by ABCG5 and ABCG8) that a locus-by-locus reading leaves as isolated signals. Second, it fitted these targets into the broader disease picture rather than displacing the established one, completing the lymphocyte transcription factor programme alongside the conventionally reported genes and embedding NR5A2 within a gut-liver-pancreas axis pertinent to PSC. Third, it elucidated the connection with clinically verified phenotypes that nearest-gene assignment does not reach: foremost the glycosylation defect COG6, whose multisystem phenotype spans the intestinal and biliary epithelium of precisely the PSC-IBD combination in which it recurs, together with the platelet and cross-autoimmune phenotypes that tie these diseases to documented haematological and lupus-related biology. Finally, it generated specific, testable hypotheses, such as the COG6 and NR5A2/sterol axes as candidate mediators of the PSC-IBD overlap, that now define clear targets for functional validation. In each case the LRR-sensitive targPred prediction did not contradict the GWAS signal but extended its interpretive reach, turning a list of associated loci into a set of mechanistic, experimentally addressable proposals.

## Author contribution

A.B. and D.D. have designed the study. A.B. has acquired funding for this work. D.D. and K.M. have designed the pipelines and performed all the analyses. D.H. has supervised targPred-related data extraction and post-proocessing. A.B. and D.D. drafted the manuscript. All authors contributed to the writing of the final manuscript. All authors read and approved the final manuscript.

## Funding

This work was conducted as a part of the “Exploring Interactions Between Regulatory Variants in Human Disease Context” project (UIP-2020-02-1623) funded by the Croatian Science Foundation. This research has also been supported by the European Regional Development Fund under grant agreement PK.1.1.10.0007 (DATACROSS). Katarina Mandić is funded by the Croatian Science Foundation project DOK-2021-02-7831.

## Code availability

Code for the analysis is available on the GitHub repository https://github.com/mlkr-rbi/UC-CD-PSC-AS_targPred.

## Notes

### Competing Interest Statement

The authors have declared no competing interest.

